# Deep Sequencing Reveals Transient Segregation of T Cell Repertoires in Splenic T Cell Zones During an Immune Response

**DOI:** 10.1101/229013

**Authors:** J. Textor, A. Fähnrich, M. Meinhardt, C. Tune, S. Klein, R. Pagel, P. König, K. Kalies, J. Westermann

**Author notes:** shared first author (J. Textor, A. Fähnrich).

## Abstract

Immunological differences between hosts, such as diverse T-cell receptor (TCR) repertoires, are widely credited for reducing the risk of pathogen spread and adaptation in a population. Within-host immunological diversity might likewise be important for robust pathogen control, but to what extent naïve TCR repertoires differ across different locations in the same host is unclear. T-cell zones (TCZs) in secondary lymphoid organs provide secluded micro-environmental niches. By harboring distinct TCRs, such niches could enhance within-host immunological diversity. On the other hand, rapid T cell migration is expected to dilute such diversity. Here, we combined tissue micro-dissection and deep sequencing of the TCR *β* chain to examine the extent to which TCR repertoires differ between TCZs in murine spleens. In the absence of antigen, we found little evidence for differences between different TCZs of the same spleen. Yet, three days after immunization with sheep red blood cells, we observed a >10-fold rise in the number of clones that appeared to localize to individual zones. Remarkably, these differences largely disappeared at 4 days after immunization, when hallmarks of an ongoing immune response were still observed. These data suggest that in the absence of antigen, any repertoire differences observed between TCZs of the same host can largely be attributed to random clone distribution. Upon antigen challenge, segregated TCR compartments appear and disappear within days. Such “transient mosaic” dynamics could be an important barrier for pathogen adaptation and spread during an immune response.

## Introduction

The evolutionary arms race between vertebrates and pathogens has led to the emergence of diversity as a core property of the adaptive immune system [1–5], enabled by sophisticated genetic mechanisms such as T-cell receptor (TCR) repertoire generation by V(D)J-recombination [5] and MHC polymorphism [3]. Between-host immunological diversity reduces the risk of pathogen spread in a population, because the huge potential diversity of the overall population (~10^15^ for human TCRs [6]) permits robust pathogen recognition on the population level. Since in humans only ~10^7^ different TCR specificities are realized per time point [7], only a very small proportion of the potentially possible TCR repertoire is present in any given individual. Thus, information about the distribution of the different TCR specificities between subjects are necessary in order to understand how infectious diseases spread and evolve within a population, which could deliver insight into how infections such as influenza [8] or HIV [9,10] could be curtailed.

Within a host, the TCR repertoire is not homogenous either. T cells are scattered across a complex eco-system consisting of secondary lymphoid organs (SLOs) such as lymph nodes, the spleen and Peyer’s patches [11]. Moreover, SLOs themselves are sub-structured into clearly separable zones that harbor distinct lymphocyte subpopulations. For instance, periarteriolar lymphoid sheaths in the spleen mainly contain T cells (T-cell zone, TCZ). It is conceivable that this microstructure fosters a diverse ‘within-host’ TCR ecosystem, with different TCZs harboring distinct TCR specificities. This might likewise make it harder for a pathogen to spread within a host, as it will trigger different local immune responses that would target a larger number of epitopes and make ‘immune escape’ more difficult. Conversely, localization of pathogens to certain specific environments could be a strategy to circumvent this effect, as with HIV which exerts most of its depletion of CD4 T cells within gut-associated lymphoid tissue [12].

However, it is unclear to what extent two TCZs indeed differ regarding their TCR repertoire. Factors that might diversify these repertoires include local homeostatic proliferation of T cells [13], differential migration of cells to specific SLOs [14,15], as well as selective local death of T cells. Homogenizing effects include T cell death (which reduces overall diversity) and, perhaps most importantly given its speed, random migration of T cells between SLOs [16]. In addition, it is not understood how an immune response affects local diversity. Does it decrease the difference in TCR diversity between the two TCZs because the same TCRs respond in both TCZs, or does it increase the difference because different TCRs are recruited into the immune response?

Recognizing and quantifying these mechanisms has also important implications for comparisons of TCR repertoires across individuals. For instance, a part of the difference between “public” clones (TCRs that are found in many different individuals) and “private” clones (not shared between individuals) found in humans [17] could be attributable to the fact that certain clones migrate more between SLOs and thus more frequently travel through the blood, from which T cells in humans are typically drawn for examination. Thus, if the exchange of TCR specificities between TCZs is slow, then the true overlap of TCR repertoires between humans could be difficult to estimate from the blood.

Here we aimed to quantify the extent to which TCR repertoires of different TCZs of the same SLO differ in mice. Specifically, we asked the following questions: (1) How do differences between two TCZs of the same murine spleen compare to those between mice? (2) How much, and how fast, do these differences change upon antigen exposure? To address these questions, we combined tissue micro-dissection with deep sequencing of the complementarity determining region 3*β* (CDR3*β*) to interrogate the TCR repertoire in murine splenic TCZs in the absence of antigen, as well as at various time-points after exposure to sheep red blood cells (SRBC), a complex non-proliferating antigen.

## Results

### Identification of CDR3*β*-Sequences from Individual Splenic TCZs

To interrogate the T-cell receptor (TCR) repertoire heterogeneity across splenic T-cell zones (TCZs) in the naïve state, we identified and isolated two separate TCZs each from serial sections of three murine spleens (Figure 1a). By determining the volume of each TCZ, we estimated that these TCZs contained about 1.9 x10^5^ T cells on average. After micro-dissection, the TCZs were subjected to high-throughput sequencing. On average, 1.6 x10^6^ reads per TCZ could be mapped to an assembled CDR3*β* gene (Figure 1b). Using the RTCR pipeline [18], we reconstructed the TCR repertoire from these reads while correcting for possible sequencing errors. Because we were primarily interested in the functional repertoire, we treat sequences with different nucleotide sequences but identical amino acid sequences as the same sequence. Therefore, throughout this study, we use the term “clone” to identify all T cells whose CDR3*β* sequence is identical on the amino acid level. This means that two cells of the same clone can still differ in CDR1 and 2, and in their alpha chain.

**Figure 1:**
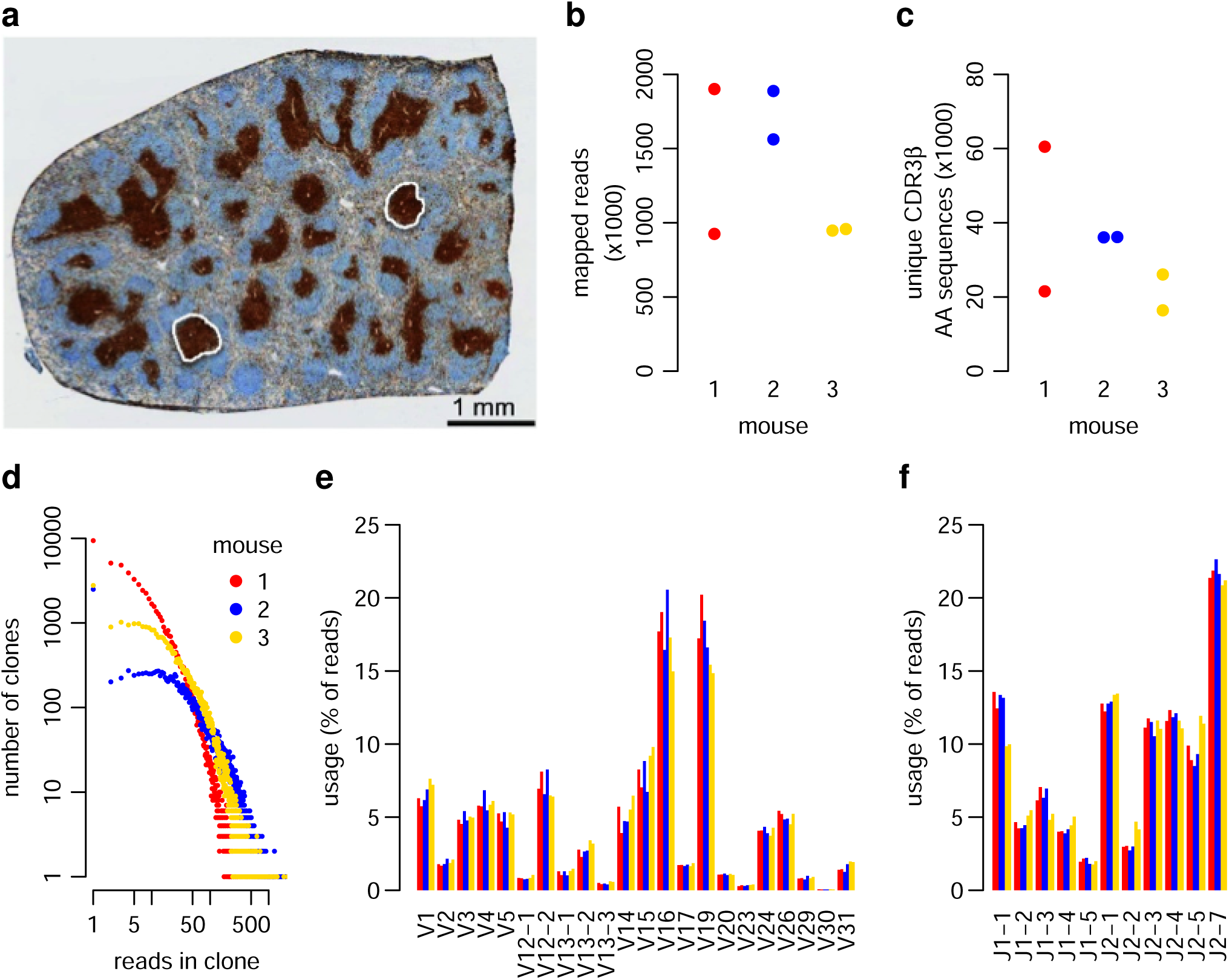
Interrogating the local repertoire in splenic T cell zones in mice. **(a)** Cross-section of a murine spleen with two separate T cell zones (TCZs) highlighted. **(b)** Number of reads mapped to the CDR3*β* region for six TCZs in three mice. **(c)** Number of unique productive CDR3*β* amino acid sequences extracted from the reads in (b). **(d)** Distribution of mapped reads per clone. **(e, f)** Overview of **(e)** V gene and **(f)** J gene usage within each sample.

The number of unique clones found per TCZ ranged from ~18,000 to ~60,000 (Figure 1c). The distribution of reads mapped to individual clones showed the expected pattern of few clones to which many reads were mapped, and many clones to which few reads were mapped, with the singleton clones representing the largest share of the reads in each sample (Figure 1d). The usage of V genes (Figure 1e) and J genes (Figure 1f) was highly consistent across the TCZ. Thus, our combination of tissue micro-dissection and deep sequencing produced CDR3*²* sequence samples with reproducible properties.

### Clonal Overlap between TCZs in the Naïve State

Repertoire comparisons between two different hosts routinely show that a small number of clones are shared, whereas a larger number are unique, i.e., only found in one host. To some extent, this large number of unique clones is simply a consequence of sampling only a small part of the underlying repertoire. However, there are also systematic differences between TCRs as to their likelihood of being generated during somatic recombination; for instance, a TCR with 3 specific nucleotide additions has a higher likelihood of being made than another one with 6 specific nucleotide additions. These differences lead to differences in average clone size, which can explain the overlap patterns between hosts to a large extent, perhaps even fully [19]. We were interested in the extent of overlap between two TCZ of the same spleen, and how this compares to the overlap between mice.

As a first attempt, we computed the symmetric overlap (Jaccard index, see Methods) between the TCZs, expressed as a percentage; for instance, a Jaccard index of 10% means that 10% of the pooled clones from two TCZ are shared (present in both TCZ). Comparing the TCZ of the same mouse, we found symmetric overlaps of 6%, 7%, and 5%, respectively, whereas the symmetric overlaps between TCZ of different mice ranged between 4% and 7% (Figure 2a). Thus, with respect to the clonal overlap, TCZs from the same mouse differed by very similar amounts as TCZs from different mice.

**Figure 2:**
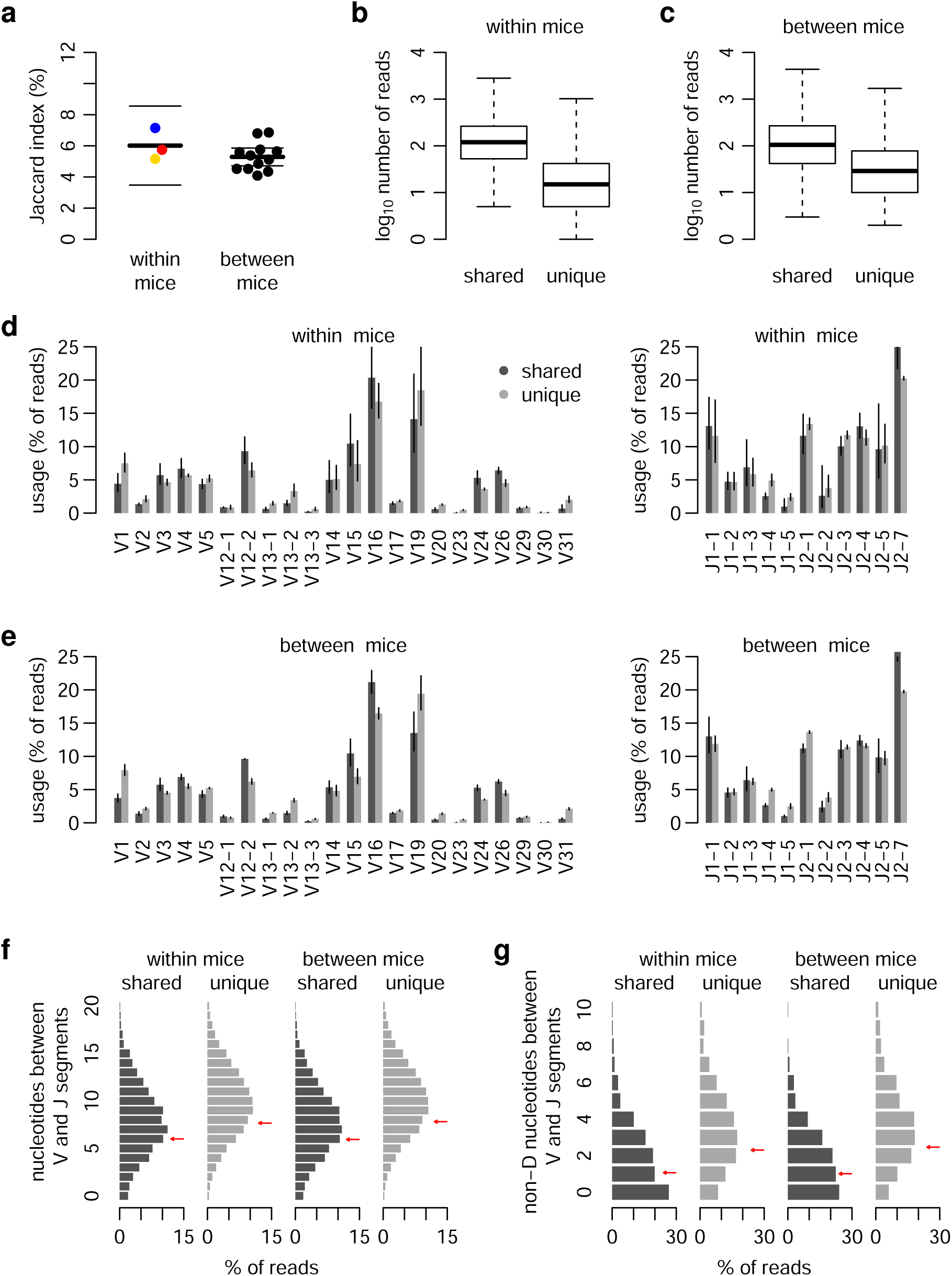
TCZ repertoire overlap is similar within and between mice, and related to precursor frequency. **(a)** Jaccard index (percentage of clones in the pooled sample occurring in both samples) for each pair of TCZs. **(b, c)** Number of reads for shared and unique clones on the within-mouse (b) and between-mouse levels (c). Boxplots show medians and interquartile ranges, whiskers show minima and maxima. **(d)** V and J gene usage for reads whose TCR sequence is shared or not shared (unique) between two TCZs of the same mouse. Error bars depict 95% confidence intervals (CIs). **(e)** V and J gene usage of reads shared or not shared between each pair of mice after pooling the reads from the two TCZs of each mouse. Error bars: 95% CIs. **(f)** Distributions of the distance in number of nucleotides between the end of the mapped V segment and the beginning of the mapped J segment. Arrows depict means. **(g)** Distribution of the number of nucleotides between V and J in each read that cannot be mapped to a D segment. Arrows depict means.

We next evaluated the basic properties of clones that were shared or unique between two TCZs of the same mouse. Both on the within-mouse (Figure 2c) and the between-mouse level (Figure 2d), we saw an expected trend in which shared clones had higher read counts than unique clones. We next evaluated usage of V and J genes. On the within-host level, no striking differences between shared and unique clones were seen (Figure 2d), except for the V31 gene, which was >2-fold more abundant in the shared clones. The V and J gene usage of shared and unique clones between hosts was likewise virtually indistinguishable (Figure 2e). Moreover, the shared clones had slightly shorter sequences (Figure 2f) and a 2-fold lower number of nucleotides that could not be mapped to a V, D, or J gene segment (Figure 2g), suggesting that they were closer to the germline. As such, these properties of shared sequences likely arise because they have a higher probability to be generated during VDJ-recombination and, as a consequence, occur more frequently in the host [19]. Overall, the within-host comparison thus gives a very similar picture to the between-host comparison.

### Statistical Model Reveals Repertoire Differences between Mice

One might expect two TCZs from the same mouse to be more similar than two TCZs from different mice, because two TCZs in the same spleen are populated from the same underlying repertoire. The absence of evidence for differences in clonal overlap between the within-host and between-host levels is therefore surprising, and could mean that such differences are absent, or could be subtle and affect only a small part of the repertoire, which would not change the overlap substantially. We therefore analyzed the repertoire overlap in a more detailed manner. A potential disadvantage of clonal overlap analysis is that it considers only presence or absence of a clone, and ignores the number of reads. While the relationship between read counts and underlying clone sizes is highly stochastic due to the inherent PCR heterogeneity [20] as well as possible differences in the amount of RNA contained in each cell, there is nevertheless an expected correlation between the number of mapped reads and the number of cells that were present in the sample. Therefore, we devised a statistical model to try and characterize the difference between TCZs in more detail, while taking into account the stochastic relationship between clone size and read count.

Intuitively, our model first assumes that all observed differences are purely due to stochastic heterogeneity in RNA content and/or the sequencing process, and then examines which individual clone differences are not well explained by this assumption using a likelihood ratio test. An example output of this model is shown in Table 1. By applying a p-value cutoff, we can dichotomize the clones into “segregated” and “non-segregated” ones, where “segregated” means that a clone appears to be substantially enriched in one of the two zones. Using the common arbitrary cutoff of 0.05, we do see more segregation between TCZs of different mice than within mice (Figure 3a). Moreover, out of the 15 possible ways to match TCZs to each other, the one that minimizes the total segregation between matched TCZs coincides with the correct matching of TCZs to mice (Figure 3b, Supporting Figure 1). Hence, our model appears capable of discriminating TCZs of different mice. We note that the amount of clones for which we find within-mice segregation at a p-value of <0.05 is lower than 10 for each mouse. Hence, this method delivers very little evidence for selective accumulation of TCRs in individual TCZs in the absence of antigen.

**Table 1:**
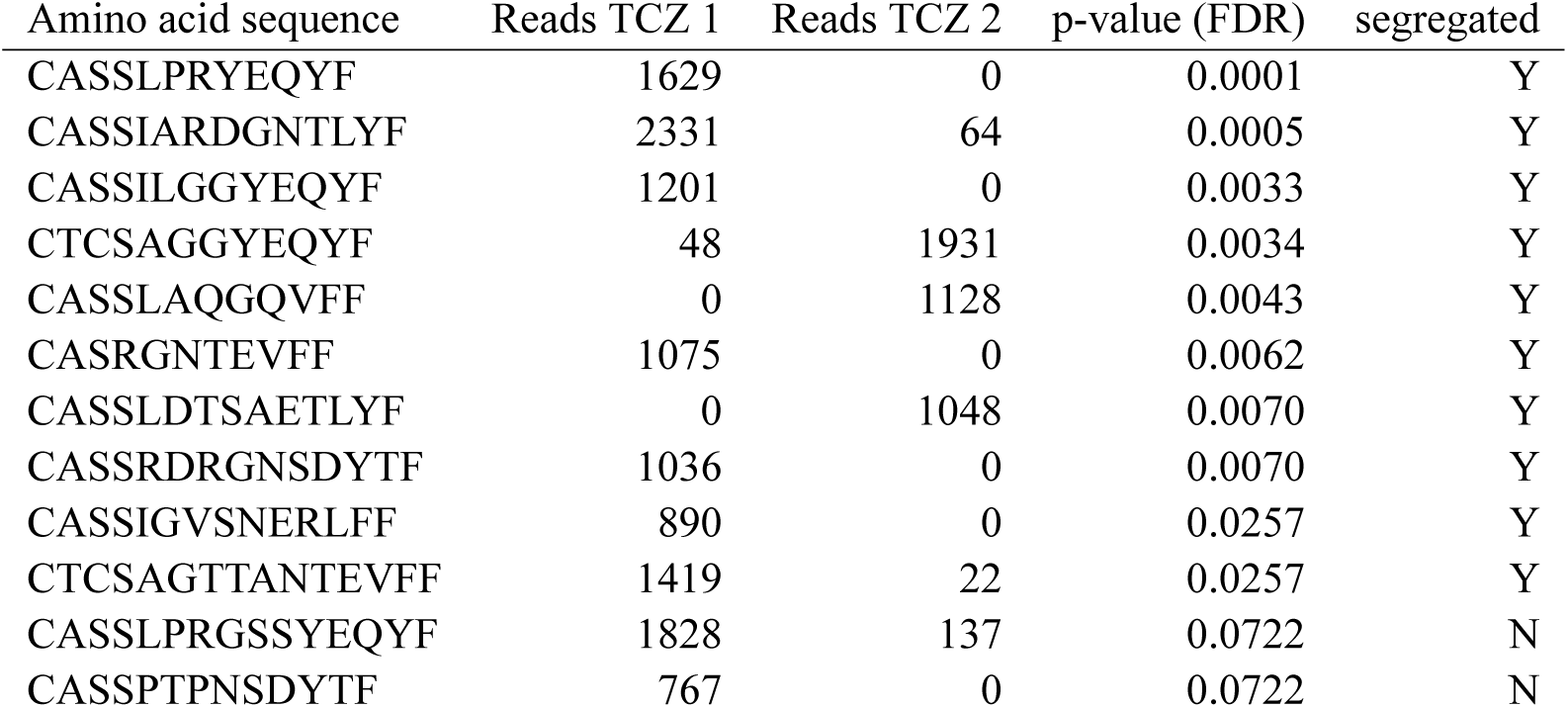
A statistical model to investigate segregation of clones within mice. This table shows an example in which the raw read counts of sequences from corresponding samples are assigned p-values for a null distribution of no difference between samples or across sequences tested using a likelihood ratio test. The p-values are then corrected for multiplicity using the FDR procedure, and dichotomized into high and low. Sequences with low p-values are called “segregated”. Note that these p-values are used here as a means to measure the divergence of read counts between samples, and not for testing a pre-specified hypothesis.

**Figure 3:**
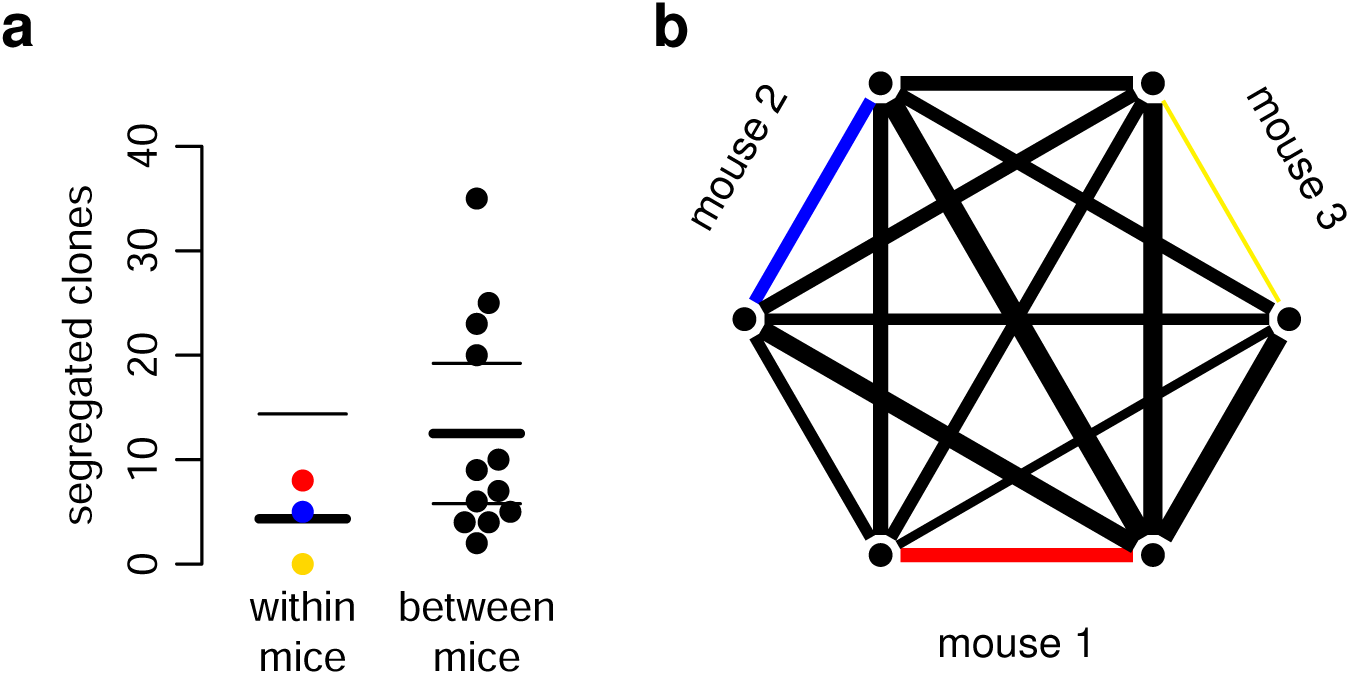
Segregation model reveals differences between mice. **(a)** Number of clones that our model classifies as “segregated” for each comparison of two TCZs of the same mouse. Lines depict mean and 95% CIs based on a t distribution. **(b)** Matching analysis in which we seek to pair TCZs to each other such that the sum of segregated clones within each pair is minimal. Line thickness is proportional to the number of segregated clones. The colored lines show the minimal matching, which corresponds to the true matching of samples to mice.

### Antigen Challenge Induces Segregation of TCZ Repertoires

To evaluate whether and how the segregation of TCR repertoires across TCZs changes during an immune response, we used sheep red blood cells (SRBCs) as antigen because a) it reaches the spleen within seconds after injection, b) no adjuvants is necessary to induce the immune response, c) it is removed from the circulation within one hour, and d) it does not proliferate [21]. This made it possible to directly and quickly induce an immune response, which is not influenced by a prolonged release of antigen and adjuvants. Thus, SRBC were injected into three mice and two splenic TCZ were collected from each mouse 3d after injection. The basic characteristics of each sample with respect to cell numbers, read counts, clone size distribution (data not shown), and VDJ status (Supporting Figure 2) were all similar to the naïve state. However, within-host TCR repertoire segregation increased strongly compared to the naïve state, as measured by a >10-fold higher number of clones that were found to be segregated (Figure 4a). Between-host TCR segregation increased by a similar amount (Figure 4b). Like in the naïve state, our model was still able to identify the correct matching of TCZs to mice (Figure 4c), suggesting that even after antigen challenge, TCZ repertoires from the same mouse remained more similar than from different mice.

**Figure 4:**
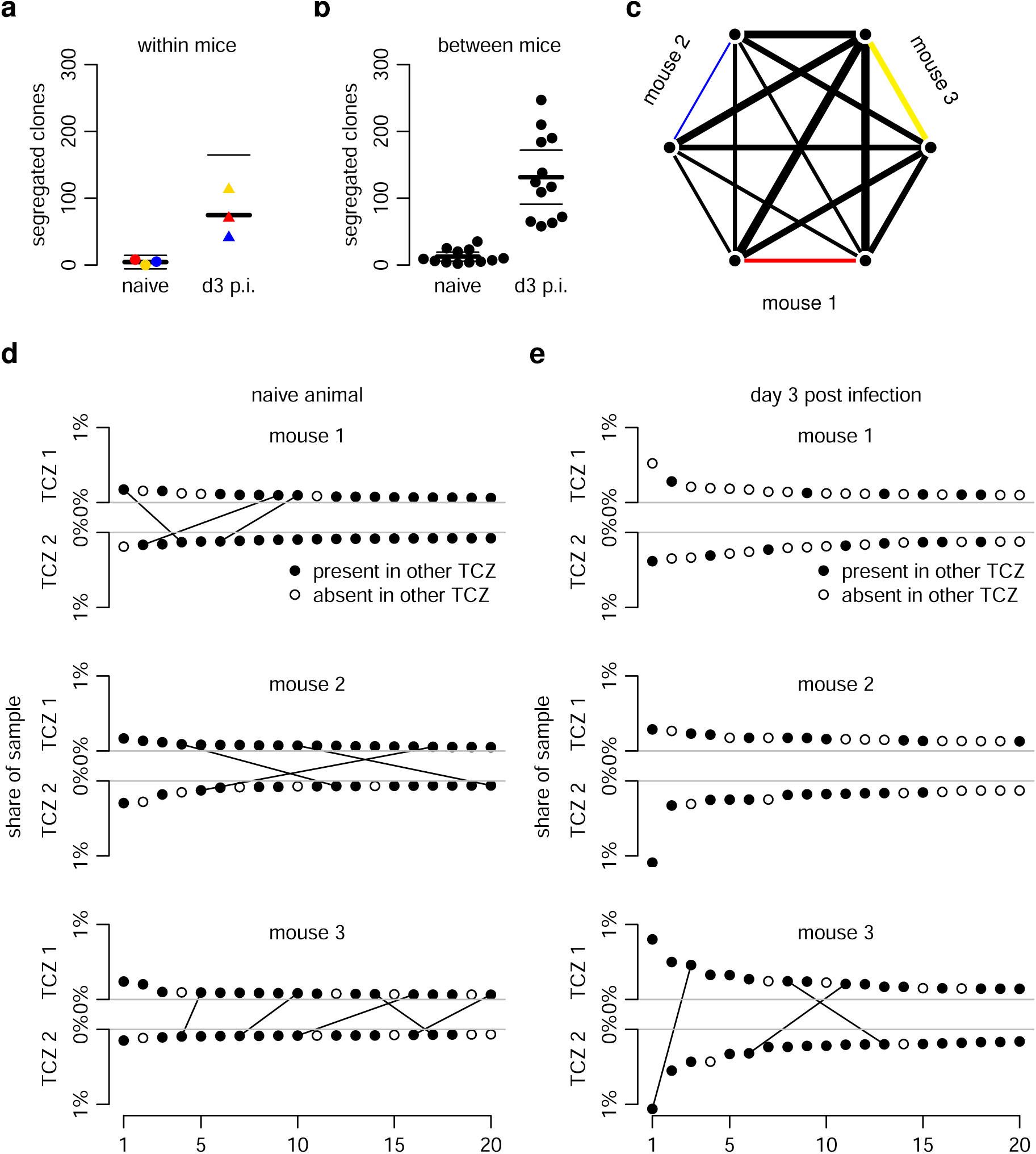
The CDR3*β* repertoire segregates within TCZs upon antigen challenge. **(a,b)** Segregation within mice increases after SRBC challenge both within (a) and across (b) mice. Lines: means and 95% CIs. **(c)** Optimal matching analysis (see Figure 3b) still correctly matches samples to mice after antigen challenge. **(d,e)** Comparison between the 20 most frequent clones in each TCZ to the other TCZ of the same mouse in the naïve state (d) or at day 3 after SRBC challenge (e) showing a decrease in the number of clones also present in the other TCZ (filled circles) after infection. Clones present in the top 20 of both TCZs are linked with a line.

To further corroborate these findings, we assessed within-host segregation in a different manner by focusing on the 20 most frequent TCR clones from each TCZ. When these were absent from the other TCZ in the same animal, we called them “segregated”. Also this method pointed to an increase in segregation 3d after antigen challenge (Figure 4d, e). Thus, our analysis indicated that antigen challenge leads to the accumulation or proliferation of different TCR clones in specific TCZs, as measured by an increase in both within-host and between-host differences between TCZs. However, TCR repertoires from TCZs in the same mouse remain more similar to each other than those from different mice.

### Local TCZ Repertoires Rapidly Desegregate During Ongoing Immune Response

To determine how long the within-host segregation of TCR repertoires persists upon antigen challenge, we injected SRBC into 3 more mice and sequenced two TCZs from each mouse 4d after antigen challenge. Surprisingly, our segregation analysis within mice (Figure 5a) and between mice (Figure 5b) as well as the top-20 overlap analysis (Supporting Figure 3) all indicated that segregation between TCZs both within and between mice had already reverted to naïve-like levels by that time. In contrast, histological analysis (Figure 5c), cell proliferation (Figure 5d), and germinal center formation (Figure 5e) all clearly showed that the immune response was still ongoing. Therefore, a likely explanation for the disappearance of segregation was egress of primed antigen-specific T cells from the TCZ, which would reduce diversity. In particular, a differential gene expression analysis using the edgeR software identified three CDR3*β* sequences whose transcription levels estimated from the blood differed between naïve and antigen-challenged mice. In all cases, the transcription levels of these CDR3*β* in the blood increased from the naïve state to d3, and increased further from d3 to d4 (Figure 5f), whereas the transcription levels in the splenic TCZs also increased from the naïve state to d3, but then remained similar or even decreased from d3 to d4. Collectively, these data show that the T-cell immune response had not yet reached its peak at d3, and was still ongoing at d4. These findings lend support to our hypothesis that egress of primed T cells from the spleen was responsible for the observed desegregation at d4.

**Figure 5:**
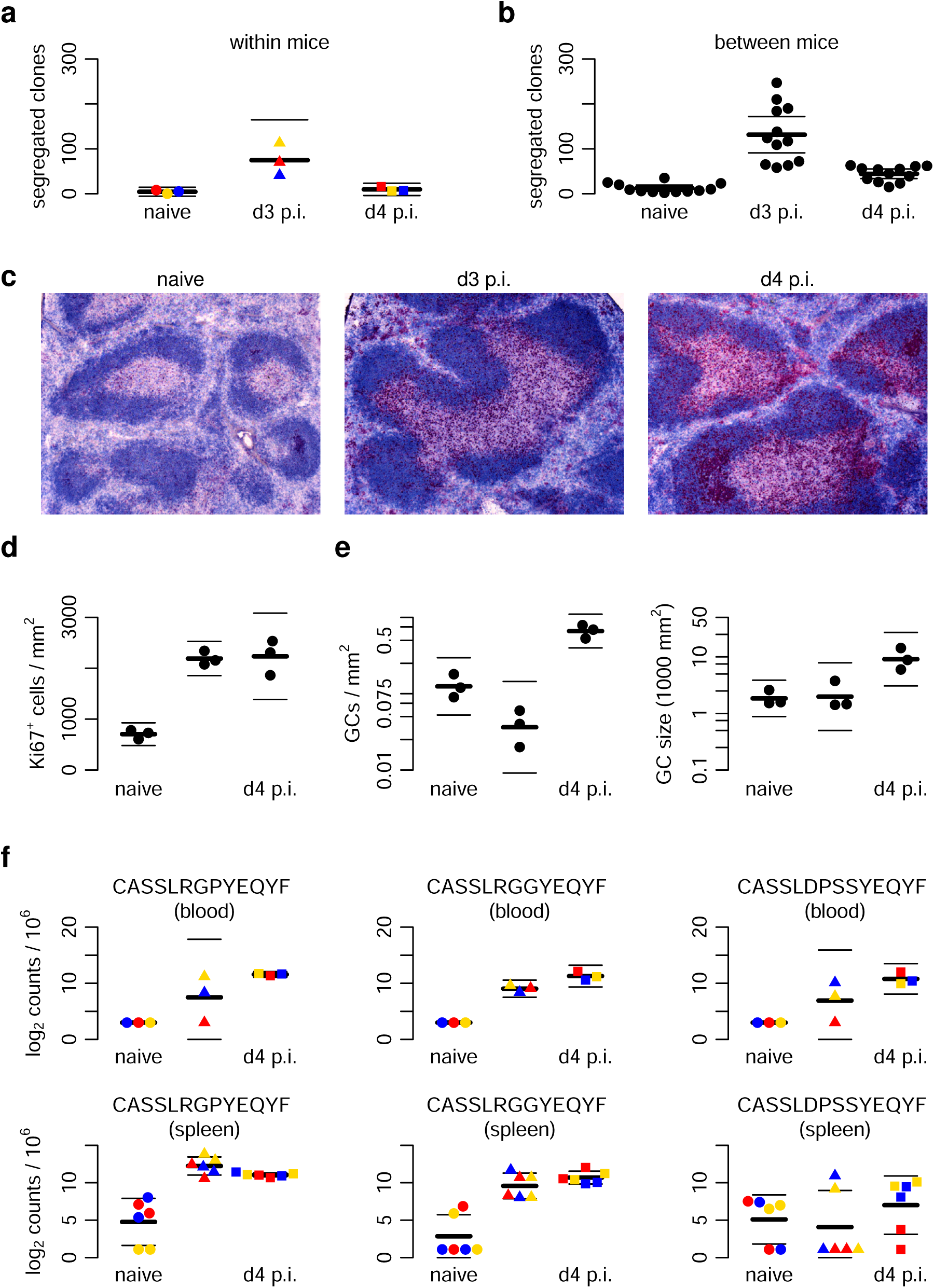
The CDR3*β* repertoire desegregates despite ongoing immune response. **(a, b)** TCZ segregation within (a) and between (b) mice reverts back to naïve-like levels at 4 days after SRBC challenge. **(c)** TCZs in a representative animal remain enlarged at d4 compared to naïve controls (B cells: blue; proliferating cells: red). **(d)** Cell proliferation in spleen remains ongoing at d4. **(e)** Germinal center formation begins at d4. **(f)** The three clones whose read counts in the blood differ most between naïve and antigen-challenged mice (top row) also remain present in splenic TCZs at d4. Throughout, lines depict means and 95% CIs.

## Discussion

Here we aimed to quantify the differences between murine splenic T-cell zones (TCZs) as well as the changes of these differences upon immunization. We found that in the naïve state, T-cell receptor (TCR) repertoires from different TCZs do differ, but these differences are smaller in magnitude compared to between-mice differences. Both within-host and between-host differences could be largely explainable by underlying heterogeneity in probability of receptor generation [22,23], leading to natural differences in clone size, combined with a simple random distribution of T cells across zones.

Upon antigen challenge, however, the situation changes dramatically and within 3d, genuinely distinct TCR micro-compartments form with differences that can no longer be explained by clone size heterogeneity and random distribution alone. Remarkably, the overlap pattern reverts back to a naïve-like state just 1d later. Thus, TCZs form what in ecology is called a “mosaic” [24]. Since most T cells constantly migrate between different SLOs as well as within individual SLOs, a “shifting mosaic” system is created [25] in which diversity could increase or decrease over time. Our findings now show that a mosaic transiently emerges in local TCZs upon antigen challenge. Our data suggests that at steady state, random migration far outpaces local proliferation and death, leading to an almost “well-mixed” situation where any diversity arises simply by chance. In contrast, after SRBC application, antigen-driven proliferation of T cells presumably increases by an extent that is sufficient to outpace intra-zonal migration, at least for a short while.

As alluded to above, shifting-mosaic dynamics may have important consequences for within-host pathogen spread and adaptation. If all TCZs of a host were to form the same immune response, a lower number of epitopes could be targeted, and a pathogen could escape the immune response through a lower number of ‘escape mutations’. Conversely, pathogens could seek to avoid immune response diversification by remaining secluded to certain areas. An even more sophisticated strategy, in which the pathogen would essentially play “cat-and-mouse” with cytotoxic T cells by appearing and then disappearing in different TCZs so quickly that the immune system cannot keep up and the virus manages to escape [26]. Our data lend empirical support to the existence of such shifting-mosaic dynamics in the T cell adaptive immune system.

Despite the enormous differences between individual TCR repertoires, there are some aspects that are consistent between individuals, sometimes to a surprising extent. One example is the phenomenon of “public” clones, which can be observed in many different individuals [17,27,28], although this, like the shared clones that we see between mice, could be simply attributable to the probability of a clone being generated and passing positive and negative selection [22,29,30]. Another example is the phenomenon of “immune-dominant” epitopes [31], in which the same epitopes of complex antigens are consistently responded to in many different individuals, a famous example being the SLYNTVATL epitope of HIV [32]. Given these effects, our finding that both intra-individual and interindividual TCZ differences increase upon immunization was not necessarily expected, since one could expect the inter-individual differences to decrease if immune responses would “converge” to a similar, “public” set of TCRs, like it has been observed on the between-host level for the hen white egg lysozyme antigen [27]. It is conceivable that such convergence may be less likely for a complex antigen like the SRBC that we used in our system. It may therefore be of interest to repeat our study using a simpler antigen or using peptide-pulsed dendritic cells to see whether TCR repertoire convergence could be observed in such a model.

Our results also have important implications for the study of clonal TCR repertoire overlap between individuals, for instance, when determining “public” and “private” clones that respond to a certain antigen. Especially when working with human subjects, the TCR repertoire is usually determined from blood samples. When there is significant within-host segregation of the TCR repertoire, the distribution of T cells in the blood may not necessarily reflect the situation in the SLOs. Our data suggest that in the absence of antigen, using blood samples to determine the clonal overlap between individuals would lead to a rather accurate estimate. However, at certain time points upon antigen challenge, the locally segregated clones – and thus a potentially important part of the immune response – would likely be missed. The extent of this potential bias would also appear to be extremely sensitive to the time point at which the measurements are taken.

In summary, our study shows that the local TCR repertoires of TCZs in the same mice rapidly segregate and de-segregate upon SRBC challenge. It remains to be determined whether this segregation is a result of purely stochastic differences in T cell localization followed by proliferation, or whether also involves clone-specific differences in cell migration or retention.

## Materials and Methods

### Mice and injections

Eight-week-old female wild type C57BL/6 mice were obtained from Charles River Breeding Laboratories, housed in the central animal facility of the University of Lübeck. All experiments were done in accordance with the German Animal Protection Law and were approved by the Animal Research Ethics Board of the Ministry of Environment (Kiel, Germany, # V312-72241.1221-1 (53-5/07). Sheep red blood cells (SRBCs; Labor Dr. Merk, Ochsenhausen, Germany) were washed and suspended in 0.9% NaCl. For immunization, 200 μl containing 10^9^ SRBC were injected into the tail vein [21]. The spleens were taken before (control), 72 h, and 96 h after injection. They were snap frozen and stored at −80°C.

### Histological analysis

Cryo-sections (thickness: 12 μm) were mounted on glass slides and stored at −80°C. To visualize the T and B cell zones (TCZs, BCZs) of the spleen, the sections were stained by immunohistochemistry using biotinylated monoclonal antibodies (TCR*β* for T cells; B220 for B cells, both BD Biosciences) as described [21]. Proliferating cells were identified by staining for Ki-67 Ag (TEC-3; DakoCytomation, Denmark) as described [33]. To estimate the number of T cells within a single TCZ, the TCZ was completely sectioned and the total area within all sections determined (2-5 x10^6^ μm^2^, n= 6 TCZ). Randomly 20 spots were chosen (670 μm^2^ in size) and the number of T cells determined revealing a Gaussian distribution with a mean T cell number of 23 (120 spots of 6 different TCZs). By dividing the total area of the TCZ by 670 and multiplying with 23 the total number of T cells per TCZ was obtained (average 1.9 x10^5^; n= 6 TCZs).

### Laser microdissection

Cryo-sections (thickness: 12 μm) were mounted on membrane covered slides (Palm Membrane Slides, PEN membrane, 1 mm; Carl Zeiss AG, Germany) and stained with toluidine blue as described [34]. To dissect splenic TCZs a pulsed UV laser was used (Palm Microbeam; Zeiss microImaging GmbH, Germany). In our hands, a TCZ area of at least 2 x 10^6^ μm^2^ is required to yield sufficient amounts of RNA for further analysis.

### Analysis of the CDR3 sequence of the TCR *β* chain

TCZs were lysed in 700μl lysis buffer (Analytik Jena, Hildesheim) as described [35]. Following RNA isolation (Analytik Jena, Hildesheim) TCR*β* chain transcripts were amplified independently in a two-step reaction according to the manufactures protocol (iRepertoire, USA, Patent No. 7,999,092,). Gene specific primers targeting each of the V and J genes were used for reverse transcription and first round PCR (One Step RT PCR Mix, Qiagen, Germany). In addition to a nested set of gene specific primers, sequencing adaptors A and B for Illumina Paired-end sequencing were added during second round PCR (Multiplex PCR Kit, Qiagen, Germany). PCR products were run on a 2% agarose gel and purified using QIAquick gel extraction kit (Qiagen, Hilden). The obtained TCRβ libraries were quantified using the PerfeCTa-NGS-quantification kit according to manufacturers protocol (Quanta, Bioscience Inc.,USA) and sequenced using the Illumina MiSeq Reagent Kit v2 300-cycle (150 paired-end read; Illumina, USA). On average, two million sequencing reads were obtained for each sample. CDR3*β* identification, clonotype clusterization and correction of sequencing errors such as removal of nonfunctional CDR3*β* sequences were performed using the RTCR pipeline [18]. Further data analysis (shared sequences, V/J gene usage, CDR3 lengths) were performed using the R platform for statistical computing. All analysis source code is available as supplementary information for this paper.

### Statistical method to determine repertoire segregation

To assess which TCR*β* sequences differed most markedly between two samples, standard differential gene expression methodology cannot be used. Therefore, we used the following simple method. First, we fitted a negative binomial distribution to the combined read counts from two samples, which assumes that each TCR*β* occurs at approximately the same frequency in each zone and that the frequencies are also approximately the same for different TCR*β*s. Then, we used the shape parameter of the negative binomial distribution and inferred the most likely location parameter for each individual pair of TCR*β* read counts using maximum likelihood. We compared these individual fits to alternative fits in which the location parameters were allowed to be different between the two samples, and computed an associated p-value from a likelihood ratio test. The full code of this method is available as Supplementary Information for reference. We emphasize that the “exploratory” p-values delivered by this method should not be seen as a basis for any kind of hypothesis test, but rather as a simple measure of a signal-to-noise ratio. Further, the method implicitly assumes that the number of segregated clones is small compared to the total number of clones, such that the variance of clone sizes samples is roughly comparable. To test the dependence of our conclusions on this assumption, we have also implemented a version of the method in which the parameters of the negative binomial distribution were fixed to the medians of the estimates obtained in all pairwise comparisons that we performed. That version gave qualitatively very similar results to those shown in this paper, and the modified model version is also available as Supplementary Information.

## Acknowledgements

We thank the members of our labs for helpful comments, and to L. Gutjahr, P. Lau, and D. Rieck for the excellent technical assistance. This project was funded by the Deutsche Forschungsgemeinschaft (DFG), grant number SFB 654, C4. JT was supported by the Dutch Cancer Society - Alpe d’HuZes foundation (project 10620).

---

**Supporting Figure 1:**
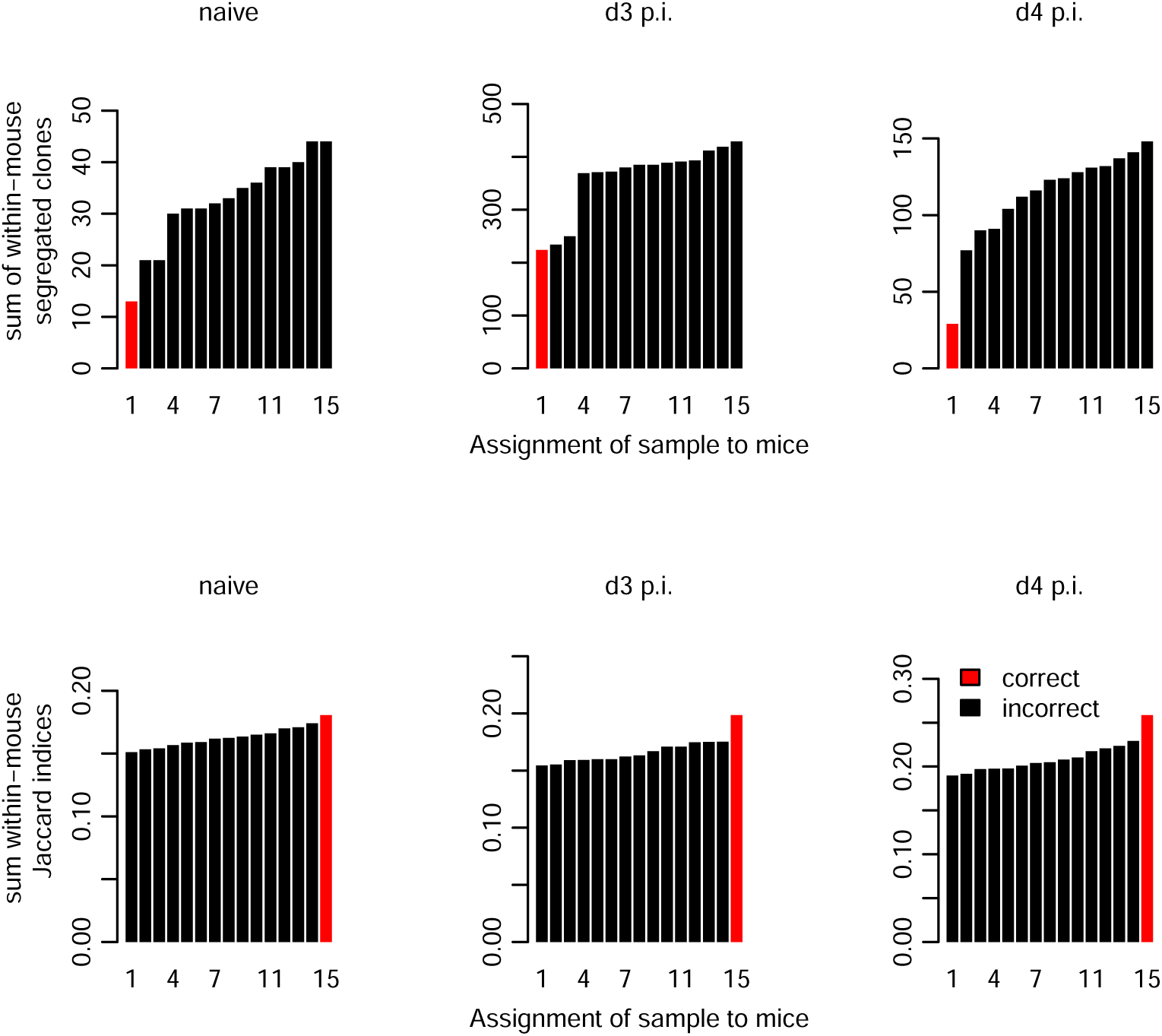
Recovering the origin of TCZ samples from mice as a validation of statistical repertoire analysis. Biologically, one expects the repertoires from two TCZs in the same mouse to be more similar than from different mice. There are 15 possible ways to match 3 mice to 2 samples each. For each of these possibilities, the plots show the sum of the number of segregated clones (top row) or Jaccard indices (bottom row) for TCZ of the same mouse. Both segregation and the Jaccard index correctly recover the correct assignment. However, the difference between the correct assignments and incorrect ones is far more pronounced for the segregation method than for the Jaccard index.

**Supporting Figure 2:**
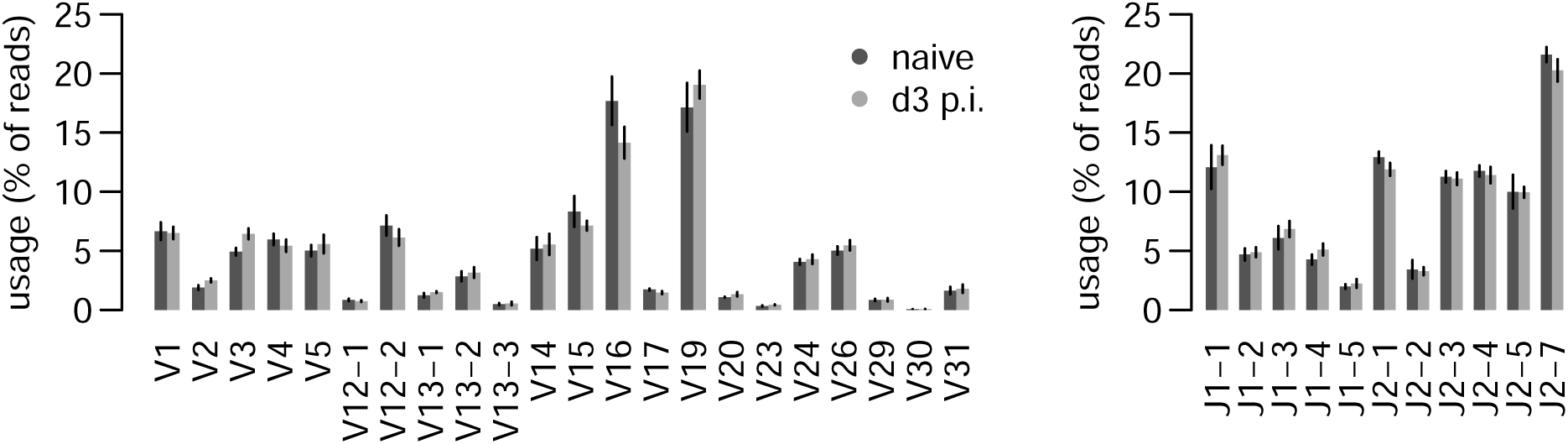
V- and J-segment usage of naïve CDR3*β* repertoires compared to 3d after SRBC infection.

**Supporting Figure 3:**
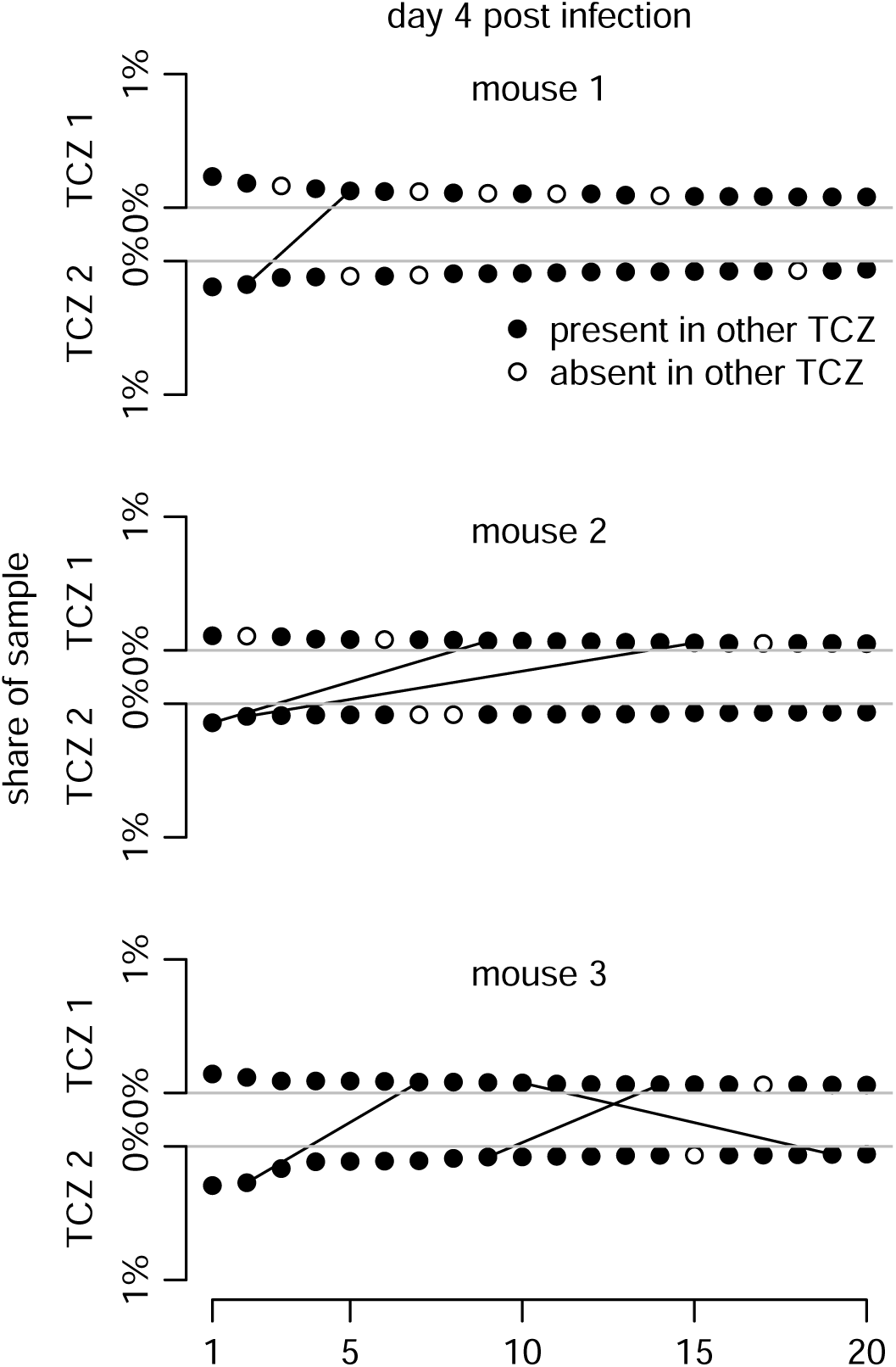
Analysis of the differences between the 20 most abundant clones in each TCZ at 4d after SRBC infection.

## References

1. Diaz M, Flajnik MF, Klinman N. Evolution and the molecular basis of somatic hypermutation of antigen receptor genes. Philos Trans R Soc Lond B Biol Sci. 2001;356: 67–72. doi:10.1098/rstb.2000.0750

2. Nikolich-Žugich J, Slifka MK, Messaoudi I. The many important facets of T-cell repertoire diversity. Nat Rev Immunol. 2004;4: 123–132. doi:10.1038/nri1292

3. Borghans JAM, Beltman JB, Boer RJD. MHC polymorphism under host-pathogen coevolution. Immunogenetics. 2004;55: 732–739. doi:10.1007/s00251-003-0630-5

4. Single RM, Martin MP, Gao X, Meyer D, Yeager M, Kidd JR, et al. Global diversity and evidence for coevolution of KIR and HLA. Nat Genet. 2007;39: 1114–1119. doi:10.1038/ng2077

5. Market E, Papavasiliou FN. V(D)J recombination and the evolution of the adaptive immune system. PLoS Biol. 2003;1: E16. doi:10.1371/journal.pbio.0000016

6. Davis MM, Bjorkman PJ. T-cell antigen receptor genes and T-cell recognition. Nature. 1988;334: 395–402. doi:10.1038/334395a0

7. Zarnitsyna V, Evavold B, Schoettle L, Blattman J, Antia R. Estimating the Diversity, Completeness, and Cross-Reactivity of the T Cell Repertoire. Front Immunol. 2013;4. doi:10.3389/fimmu.2013.00485

8. Woolthuis RG, Dorp CH van, Keçmir C, Boer RJ de, Boven M van. Long-term adaptation of the influenza A virus by escaping cytotoxic T-cell recognition. Sci Rep. 2016;6: 33334. doi:10.1038/srep33334

9. Walker BD, Korber BT. Immune control of HIV: the obstacles of HLA and viral diversity. Nat Immunol. 2001;2: 473–475. doi:10.1038/88656

10. Korber B, Gaschen B, Yusim K, Thakallapally R, Kesmir C, Detours V. Evolutionary and immunological implications of contemporary HIV-1 variation. Br Med Bull. 2001;58: 19–42. doi:10.1093/bmb/58.1.19

11. Ruddle NH, Akirav EM. Secondary Lymphoid Organs: Responding to Genetic and Environmental Cues in Ontogeny and the Immune Response. J Immunol. 2009;183: 2205–2212. doi:10.4049/jimmunol.0804324

12. Brenchley JM, Schacker TW, Ruff LE, Price DA, Taylor JH, Beilman GJ, et al. CD4+ T cell depletion during all stages of HIV disease occurs predominantly in the gastrointestinal tract. J Exp Med. 2004;200: 749–759. doi:10.1084/jem.20040874

13. Surh CD, Sprent J. Homeostatic T Cell Proliferation. J Exp Med. 2000;192: F9–F14. doi:10.1084/jem.192.4.F9

14. Mueller SN, Gebhardt T, Carbone FR, Heath WR. Memory T cell subsets, migration patterns, and tissue residence. Annu Rev Immunol. 2013;31: 137–161. doi:10.1146/annurev-immunol-032712-095954

15. Steinert EM, Schenkel JM, Fraser KA, Beura LK, Manlove LS, Igyártó BZ, et al. Quantifying Memory CD8 T Cells Reveals Regionalization of Immunosurveillance. Cell. 2015;161: 737–749. doi:10.1016/j.cell.2015.03.031

16. Textor J, Henrickson SE, Mandl JN, Andrian UH von, Westermann J, Boer RJ de, et al. Random Migration and Signal Integration Promote Rapid and Robust T Cell Recruitment. PLoS Comput Biol. 2014;10: e1003752.

17. Venturi V, Price DA, Douek DC, Davenport MP. The molecular basis for public T-cell responses? Nat Rev Immunol. 2008;8: 231–238. doi:10.1038/nri2260

18. Gerritsen B, Pandit A, Andeweg AC, de Boer RJ. RTCR: a pipeline for complete and accurate recovery of T cell repertoires from high throughput sequencing data. Bioinforma Oxf Engl. 2016;32: 3098–3106. doi:10.1093/bioinformatics/btw339

19. Murugan A, Mora T, Walczak AM, Callan CG. Statistical inference of the generation probability of T-cell receptors from sequence repertoires. Proc Natl Acad Sci. 2012;109: 16161–16166. doi:10.1073/pnas.1212755109

20. Best K, Oakes T, Heather JM, Shawe-Taylor J, Chain B. Computational analysis of stochastic heterogeneity in PCR amplification efficiency revealed by single molecule barcoding. Sci Rep. 2015;5: 14629. doi:10.1038/srep14629

21. Stamm C, Barthelmann J, Kunz N, Toellner K-M, Westermann J, Kalies K. Dose-dependent induction of murine Th1/Th2 responses to sheep red blood cells occurs in two steps: antigen presentation during second encounter is decisive. PloS One. 2013;8: e67746. doi:10.1371/journal.pone.0067746

22. Elhanati Y, Murugan A, Callan CG, Mora T, Walczak AM. Quantifying selection in immune receptor repertoires. Proc Natl Acad Sci. 2014;111: 9875–9880. doi:10.1073/pnas.1409572111

23. Elhanati Y, Marcou Q, Mora T, Walczak AM. repgenHMM: a dynamic programming tool to infer the rules of immune receptor generation from sequence data. Bioinformatics. 2016;btw112. doi:10.1093/bioinformatics/btw112

24. Singer MC, McBride CS. Geographic mosaics of species’ association: a definition and an example driven by plant-insect phenological synchrony. Ecology. 2012;93: 2658–2673. doi:10.1890/11-2078.1

25. Bormann FH, Likens GE. Pattern and Process in a Forested Ecosystem. New York: Springer-Verlag. 1979. New York: Springer; 1979.

26. Lythgoe KA, Blanquart F, Pellis L, Fraser C. Large Variations in HIV-1 Viral Load Explained by Shifting-Mosaic Metapopulation Dynamics. PLOS Biol. 2016;14: e1002567. doi:10.1371/journal.pbio.1002567

27. Cibotti R, Cabaniols JP, Pannetier C, Delarbre C, Vergnon I, Kanellopoulos JM, et al. Public and private V beta T cell receptor repertoires against hen egg white lysozyme (HEL) in nontransgenic versus HEL transgenic mice. J Exp Med. 1994;180: 861–872. doi:10.1084/jem.180.3.861

28. Venturi V, Kedzierska K, Price DA, Doherty PC, Douek DC, Turner SJ, et al. Sharing of T cell receptors in antigen-specific responses is driven by convergent recombination. Proc Natl Acad Sci. 2006;103: 18691–18696. doi:10.1073/pnas.0608907103

29. Quigley MF, Greenaway HY, Venturi V, Lindsay R, Quinn KM, Seder RA, et al. Convergent recombination shapes the clonotypic landscape of the naive T-cell repertoire. Proc Natl Acad Sci U S A. 2010;107: 19414–19419. doi:10.1073/pnas.1010586107

30. Li H, Ye C, Ji G, Wu X, Xiang Z, Li Y, et al. Recombinatorial biases and convergent recombination determine interindividual TCR*β* sharing in murine thymocytes. J Immunol Baltim Md 1950. 2012;189: 2404–2413. doi:10.4049/jimmunol.1102087

31. Yewdell JW. Confronting complexity: real-world immunodominance in antiviral CD8+ T cell responses. Immunity. 2006;25: 533–543. doi:10.1016/j.immuni.2006.09.005

32. Iversen AKN, Stewart-Jones G, Learn GH, Christie N, Sylvester-Hviid C, Armitage AE, et al. Conflicting selective forces affect T cell receptor contacts in an immunodominant human immunodeficiency virus epitope. Nat Immunol. 2006;7: 179–189. doi:10.1038/ni1298

33. Barthelmann J, Nietsch J, Blessenohl M, Laskay T, van Zandbergen G, Westermann J, et al. The protective Th1 response in mice is induced in the T-cell zone only three weeks after infection with Leishmania major and not during early T-cell activation. Med Microbiol Immunol (Berl). 2012;201: 25–35. doi:10.1007/s00430-011-0201-6

34. Kalies K, Blessenohl M, Nietsch J, Westermann J. T Cell Zones of Lymphoid Organs Constitutively Express Th1 Cytokine mRNA: Specific Changes during the Early Phase of an Immune Response. J Immunol. 2006;176: 741–749. doi:10.4049/jimmunol.176.2.741

35. Li K-P, Fähnrich A, Roy E, Cuda CM, Grimes HL, Perlman HR, et al. Temporal Expression of Bim Limits the Development of Agonist-Selected Thymocytes and Skews Their TCR*β* Repertoire. J Immunol Baltim Md 1950. 2017;198: 257–269. doi:10.4049/jimmunol.1601200

